# Block-Champagne: A novel Bayesian framework for Imaging E/MEG Source

**DOI:** 10.1101/2025.02.17.638774

**Authors:** Zhao Feng, Cuntai Guan, Yu Sun

## Abstract

Estimating the extents of E/MEG source activities is vital for exploring brain dynamics at high spatiotemporal resolution. However, conventional ESI models exclusively overemphasize on the accurate estimation of the source locations and struggle to reconstruct the extents due to the ill-posed nature of the problem, which requires effective integration of biophysical constraints and proficiency in signal processing methods. In this study, a novel Bayesian framework – Block-Champagne is introduced to provide accurate estimation of extended sources (i.e., both locations and extents). Specifically, a block-sparsity constraint is employed to model the local homogeneity of sources, which can be updated automatically to reconstruct source with arbitrary extents. Furthermore, prior constraints from other modalities, such as fMRI maps, can be incorporated to model interactions between remote sources to further enhance source reconstruction. The performance of Block-Champagne was quantitatively evaluated through simulation experiments, demonstrating its overall superiority under various complex scenarios (i.e., SNR, extent size, & number of extents) compared to benchmark algorithms (including LORETA, EBI-Convex, ts-Cham, L21-Sissy, & SI-STBF). Moreover, validation results obtained from deep-brain stimulation EEG, epilepsy data, and face-processing multi-modal data further proved the practical feasibility of Block-Champagne. In conclusion, our study reveals the superiority of the proposed Block-Champagne in accurate reconstruction of extended source, positioning Block-Champagne as a highly promising tool for realistic applications where source locations and extents are of equivalent importance.

## I. Introduction

For the unique advantage of non-invasive monitoring brain dynamics with high spatio-temporal resolution, electrophysiological source imaging (ESI) has attracted continuously growing interests in the field of neuroscience and clinical applications [1]. Heuristically, ESI techniques aim to reconstruct the underlying sources that generate the recorded electroand magnetoencephalography (E/MEG) signals, with particular emphasis on accurate estimation of the source locations and extents. Among the ESI models, the distributed model, which assumes the sources as current density spread over the cortex surface, has shown promising and neurophysiological plausible results [2]. Then, the ESI problem could be equivalently derived as to identify the most accurate current distributions that account for the observed E/MEG signals. However, the problem is inherently ill-posed, as numerous sources configurations could yield the same measurements [3]. The prior constraints (i.e., from biological and/or mathematical perspectives) are therefore essential to be incorporated in the ESI model that lead to source characteristics aligning with the biophysiological properties of neural activity. properties of neural activity.

From biological perspective, the E/MEG signals are assumed to originate from spatially extended and locally homogeneous sources, since convergent findings show that a large number of neural cells need to activate synchronously to produce detectable signals [4], [5]. As such, the solutions to ESI problem can be modeled using extended source patches, where the activities from adjacent sources are highly correlated. Conventional approaches have incorporated various mathematical constraints to solve the problem. For instance, the low resolution brain electromagnetic tomography (LORETA) [6] employs Laplacian operator within the *L*_2_ regularization term to enforce homogeneous activities between adjacent sources. Such *L*_2_ norm-based methods, often known as minimum-norm family, typically produce spatially blurred solutions. To accurately identify the location of sources, methods based on sparse constraints have been proposed, including iteratively reweighted approaches like FOCUSS [7], *L*_*p*_-norm (0 *< p* ≤ 1) regularization [8], and sparse Bayesian learning methods [9]. One major limitation of these sparse constraint techniques is that the estimated sources are over-focal. These conventional spatially-constrained models provide either blurred results without clear edges between sources or over-focal results without information of source extents. However, convergent findings have showed that source extents are of equivalent importance as accurate estimation of source location [10]. The inability to reliably reconstruct the source extents would limit the wide applications of ESI.

According to the functional specialization hypothesis, different brain regions are specialized for distinct functions [11]. Therefore, the ESI results should exhibit structural sparsity, that is, source activities are spatially clustered within each source patch and with only a few patches being activated in response to a given stimulus, further highlighting the equivalent importance of source locations and extents. To address this, advanced ESI methods have adopted various strategies. For example, the cortex parcellation approach segments the cortex into distinct parcels as SBFs, and ESI algorithms then determine the relevant SBFs to reconstruct source extents [12], [13]. Most recently, Li et al. proposed a two-stage champagne method, where the potential locations of sources were firstly identified followed by the construction and best-fitting of SBFs that represent various extents around the local sources through a champagne approach [14]. A key challenge with these methods is to construct SBFs that accurately represent the shape and extent of the underlying sources. Another widely-explored approach is to employ constraints in multiple domains, e.g., incorporating sparse constraint in gradient domain. Studies have reported that such total variation model could generate clear edges between source and background activities, thus contributing to the reconstruction of source extents [15], [16], [17]. Of note, these methods may yield suboptimal solutions when the complex algorithm hyper-parameters were not well tuned [18]. Hence, the specialized knowledge needed for the implementation of these two strategies may limit the wide application.

Another practical solution to improve the reconstruction of source locations and extents is to integrate information from other modalities [19], [20]. Among them, functional magnetic resonance imaging (fMRI) is particularly attractive for its common origins with E/MEG (although not exactly the same) [1], the feasibility to collect fMRI and E/MEG concurrently, and the high spatial resolution compared to E/MEG. One common approach is the hard-threshold strategy, where the prior source covariance is constructed based on the fMRI maps [21], or the SBFs derived from fMRI decomposition [22]. This approach forces the obtained results to be exactly consistent with fMRI maps, which could be incorrect due to the mismatch between the origins of fMRI and E/MEG [23]. Another approach is the soft-threshold strategy, where higher probabilities of activation from fMRI maps are assigned to the corresponding sources. Studies have incorporate the fMRI-derived covariance into the regularization terms, thereby reducing penalization for sources indicated by fMRI maps [24], [25]. Nevertheless, both approaches rely on fixed prior constraints solely derived from other modalities, neglecting the correspondence with the recorded E/MEG signals. More effective strategies are therefore needed to better balance the information from fMRI priors with the evidence from E/MEG data.

In this work, we propose a novel Bayesian framework − Block-Champagne (abbreviated to Blk-Cham hereafter) to reconstruct the source locations and extents through integrating spatial information from local and/or remote scale. Specifically, Blk-Cham employs block-sparsity constraints to model the local homogeneity of each source and its first-order adjacent neighbors as a single block [26]. The blocks can be combined to reconstruct arbitrary source extents in a datadriven manner. Besides, the proposed Blk-Cham allows for incorporation of activation information supported by physiological evidence or data from other modalities. In fact, inclusion of functional synchronize/desynchronize between distant brain regions could benefit in-depth understanding of the underlying source dynamics [27]. To this end, we model the dynamics of each source as a combination of its own activity and that of remote regions [28] to allow Blk-Cham capturing interactions indicated by the prior constraints. The included constraints are updated to balance the fit-to-data and prior knowledge. We have quantitatively evaluated the performance of Blk-Cham through simulated data under various conditions, which reveals the superiority of Blk-Cham through comparing with five widely-used/state-of-the-art benchmark methods. Additionally, we have also demonstrated the application of Blk-Cham to real EEG data.

## II. Methods

### A. Probabilistic generative model

According to the augmented model [29], the scalp E/MEG can be modeled as:

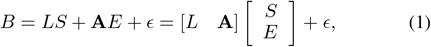

The variables in the augmented model are listed in Table I. The formulations and the derivation of the Blk-Cham model are provided below.

**TABLE 1.**
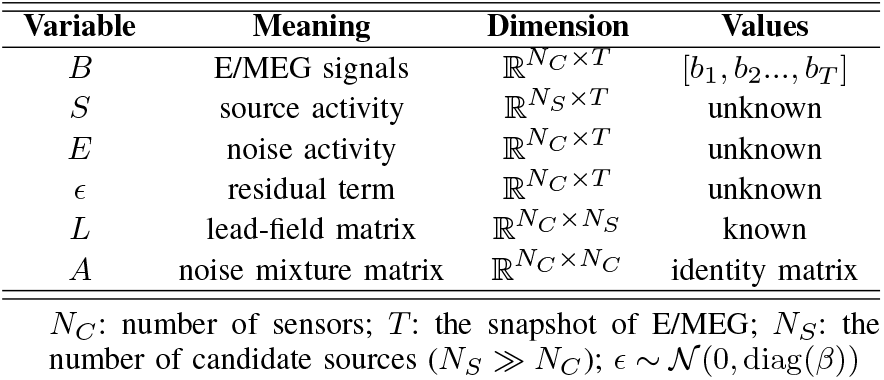
Variables in the augmented model.

#### 1) Local information

The source *S* is firstly decomposed as:

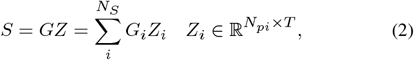

where 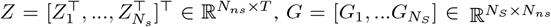 is a zero matrix except that the part from *i*^*th*^ row to (*i* + *N*_*pi*_ − 1)^*th*^ row is replaced by identity matrix,. The local homogeneity of source activities is modeled by:

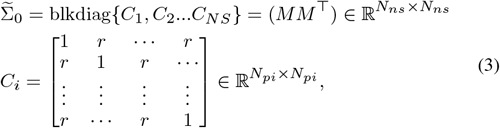

where *C*_*i*_ is the covariance for *i*^*th*^ dipole and its first-order neighbors, *N*_*pi*_ = 1+ (no. of neighbors of dipole *i*) and 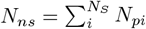. The diagonal elements of *C*_*i*_ are set to be 1 and others are *r* (*r* ≈ 1 is the intra-block correlation). The block covariance in Eq. 3 indicates that *Z* can be decomposed as:

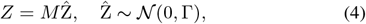

then 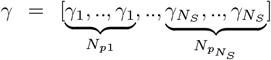 is a vector of nonnegative hyper-parameters and Γ = diag(*γ*) represents prior hyperparameters for both source and noise activity.

#### 2) Remote information

We relax the part of Γ corresponding to connectome map to be a symmetric matrix, where the off-diagonal elements represent connections between brain regions within the same connectome map (e.g., obtained from fMRI). The Γ in Eq. 4 is reformulated as:

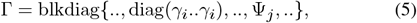

where we keep the *γ* of dipoles outside the connectome map to be diagonal, i.e., to model the local homogeneity only. On the other hand, we relax the part of dipoles inside the map (denote as Ψ) to be a symmetric matrix, i.e., to model source activities caused by local correlation and functional connectome relationship.

Taken together, the source imaging model is expressed as:

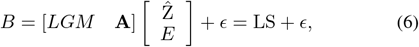

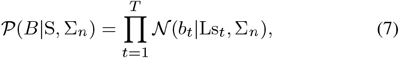

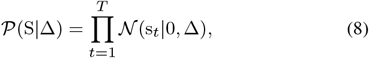

and 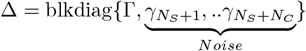.

The posterior distribution S, noise covariance Σ_*n*_ and hyperparameters *M*, Γ can be automatically learned from data. The graphic illustration of Blk-Cham is presented in Fig. 1.

**Fig. 1.**
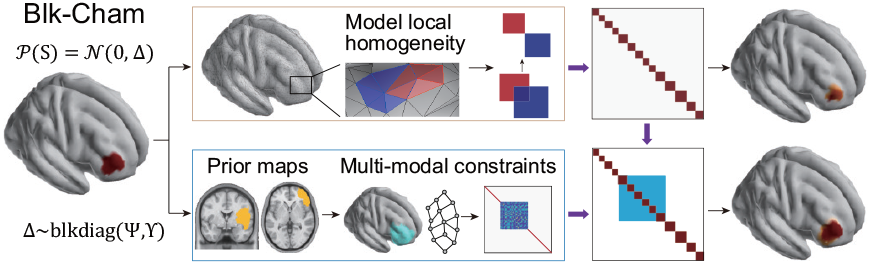
Illustration of the proposed Blk-Cham. The method models each dipole and its adjacent neighbors as one block to promote local homogeneity. The blocks can be overlapped to reconstruct arbitrary source extents. Moreover, if additional information pertaining to source activation is available from other modality, e.g., fMRI, the methods can incorporate the prior constraints to enhance performance of source estimation.

### B. Solution to source imaging model

According to the Bayes’ rule *p*(S | *B*, Σ_*n*_, *M*, Γ) ∝ *p*(S | Γ)*p*(*B*| S, *M*, Σ_*n*_), the mean of posterior distribution can be obtained while keeping other parameters fixed:

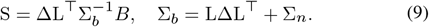

The hyperparameters can be estimated by minimizing the cost function:

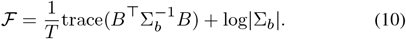

The update rule for hyperparameters Γ is derived from convex analysis of log|Σ_*b*_|. Since the log|Σ_*b*_| is a concave function with respect to ∆, the cost function can be expressed as a minimum over upper-bounding hyperplanes [9]:

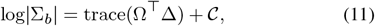

where Ω is a matrix of auxiliary variables and 𝒞 is the term that irrelevant to Γ. Similarly, trace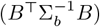 can also be expressed as:

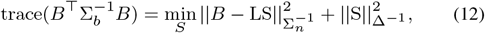

thus we get the cost function :

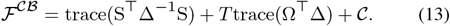

The updating rule is obtained through taking the derivative with respect to ∆, and calculating a hyperplane which is tangential to log |Σ_*b*_|. Notably, the part of ∆ (i.e., Ψ) corresponding to dipoles that locate in connectome map is a symmetric matrix, otherwise ∆ is a diagonal matrix. The update of Ψ is:

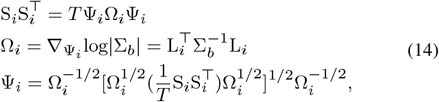

and the update of diagonal part is:

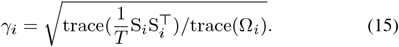

In practice, it is reasonable to assume fixed values of *C*_*i*_, i.e., adjacent dipoles share high correlation activities with the same intra-block correlation coefficient *r*≈ 1 for all dipoles [26]. Besides, the Σ_*n*_ of residual term can also set to be a small value, which guarantees the stability of the algorithm [29].

In this section, we also provide the update rules for *C*_*i*_ and Σ_*n*_ in a data-driven manner, which provide a convenient framework for automatic source estimation. The update of Σ_*n*_ = diag(*β*) is:

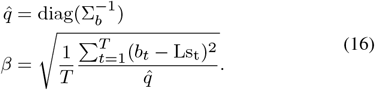

To learn the local homogeneity of source, we can take the derivation of ℱwith respect to each 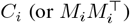. To avoid overfitting, we fix all the *C*_*i*_ to share the same form as Eq. 3 and have the same correlation *r*, thus we only need to learn the *r* from data. Without loss of generality, the *r* is updated based on the diagonal part of prior covariance *F*. By reformulating Eq. 6 as:

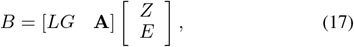

we find that the part related to *C*_*i*_ is 𝒫 (*Z*_*i*_ 0, *γ*_*i*_*C*_*i*_). Thus, the *C*_*i*_ can be updated in an expectation-maximization manner:

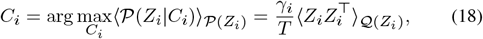

where 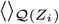 denotes expectation with respect to posterior mean of *Z*_*i*_. Once the blocks have been updated, we calculate the averages of the elements along the main diagonal and main sub-diagonal of 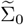, i.e., *m*_0_ and *m*_1_, and update *r* as:

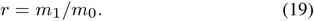

The framework of Blk-Cham is shown in Algorithm 1. The code and example data are available at https://github.com/ZJUNeural/BlockChampagne.

#### Algorithm 1 The Blk-Cham framework

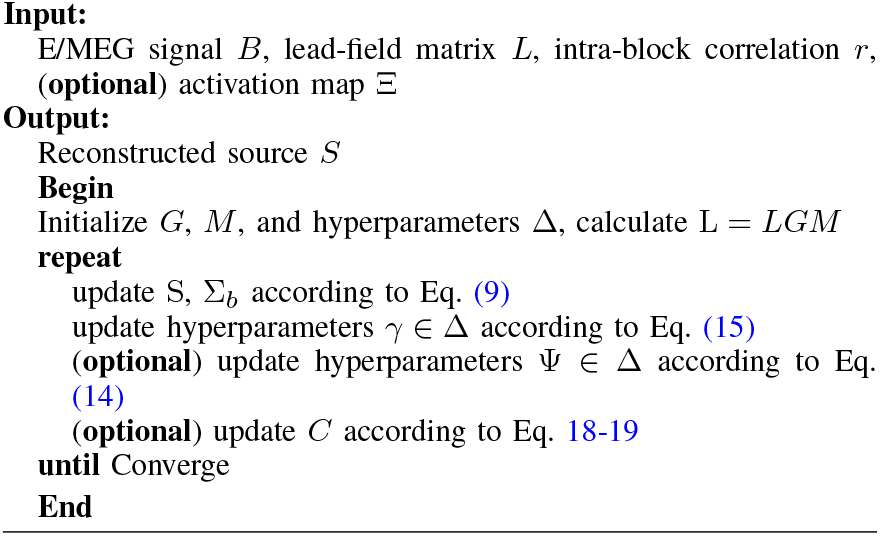

## III. Performance evaluation

The performance of the proposed Blk-Cham is evaluated using both simulated and realistic data. For the simulation experiments where ground truth could be obtained, quantitative assessments of Blk-Cham were conducted and compared with five benchmark ESI models under various conditions using four well-established evaluation metrics (i.e., AUC, DLE, MSE, and CORR). To further validate the practicality and effectiveness of Blk-Cham in real-world scenarios, we apply the method to three publicly available datasets.

### A. Simulation protocol

To generate simulated source activities, we use the standard anatomical template − ICBM152 anatomy [30] to obtain the cortical surface. The cortex is first downsampled into 4747 dipoles, with the orientation of each dipole fixed to be perpendicular to the cortical surface. We use 53 channels to obtain the simulated scalp signals, and compute the lead-field matrix (53 × 4747) using the Boundary Element Method (BEM) implemented in OpenMEEG (BrainStorm toolbox [31]). For each simulation, we randomly select seed voxels and include the adjacent dipoles of each seed voxel to obtain spatially extended sources. The temporal dynamics of source activity are generated based on event-related spectral perturbations (ERSP) model using SEREEGA toolbox [32], with Gaussian noise are added to achieve the desired signal-to-noise ratio (SNR). The SNR is defined as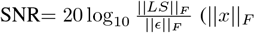 denotes Frobenius norm of. For each experiment, the metrics are calculated from 50 Monte-Carlo simulations. We quantitatively evaluate the influence of SNR, source areas and number of sources on the reconstruction performance under the following conditions, where the performance metrics are calculated:

- SNR: two source patches are activated to an extent of 5 cm^2^ under different SNR levels (i.e., −5, 0, 5, 10 dB).
- Source area: two source patches are generated with different extents (i.e., 1, 5, 10, 15 cm^2^). The SNR is set to 5 dB.
- Number of source patches: various numbers of source patches are generated simultaneously (i.e., 1−4 patches).

The source extents are around 10 cm^2^ and the SNR is set to 5 dB.

Of note, we assume no baseline information is available and set the covariance to an identical matrix in the simulation experiments.

### B. Real data validation

#### 1) Localize-MI dataset

The Localize-MI dataset is an openaccess dataset that includes 256-channel EEG data recorded from 7 subjects with stereotactically implanted electrodes, available at ^1^. The EEG data are collected simultaneously with intracortical stimulation, wherein an electrical artifact is produced and measured by scalp EEG. Therefore, the stimulation sites can serve as the ground truth of electrical currents. The EEG data are processed according to [33]. The dataset also provides cortex surface (8196 dipoles) and leadfield matrix of each subject obtained from individual sMRI. The data of subject #1 and subject #7 are used in this section. The EEG from [-0.1, 0] s prior to stimulation to calculate noise covariance for whitening, and reconstruct the source of simulated signals from [0, 0.1] s (where 0 s correspond to the onset of stimulation) using the proposed Blk-Cham. For comparison purpose, we also provide the reconstruction results of 4 benchmarks: LORETA, EBI-Convex, ts-Cham, and BESTIES.

#### 2) Epilepsy dataset

The Epilepsy dataset is an open-access dataset that contains EEG data recorded from a epilepsy patient, available at ^2^. The 27-channel EEG data consists of 58 interictal spikes, which are averaged for source localization following EEG pre-processing. The maximum of spike is recorded at FC1 channel. The individual head model is constructed using pre-surgical MRI data, and the cortex is segmented and downsampled into 6006 dipoles. The detailed information regarding data processing can be found in BrainStorm tutorial, and further analysis of this dataset as well as the surgical resection outcomes have been reported in [34]. In this study, we employ Blk-Cham to reconstruct the source of averaged spikes, with results from LORETA, BESTIES, and ts-Cham also provided for comparison.

#### 3) Face-processing dataset

The face-processing dataset is an open-access dataset that contains multi-modal neuroimaging data from 16 subjects undergoing face-processing tasks [35]. We use the concurrent EEG and fMRI, as well as sMRI data from subject #15 following the guidelines outlined in the SPM12 manual. The event-related potential (ERP) data are averaged from EEG data (70 channels, 295 trials) corresponding to the face presentations. The pre-stimulus data from [-100, 0] ms relative to the stimulus onset are used to baseline correct the ERP. In this study, we use the ERP data collected when subjects response to the familiar face stimulus. Individual sMRI data are used to generate cortex surface and leadfield matrix (8196 sources) in SPM12 based on BEM model. The task-related BOLD responses are calculated from fMRI data for comparison purpose. Detailed information regarding data processing, along with Matlab scripts for the processing procedures, are provided in [35] and Chapter 43 of SPM12 Manual. The proposed Blk-Cham, along with LORETA and BESTIES, is applied to reconstruct the underlying source of ERP activities. Of note, we also present the ESI results of Blk-Cham that incorporates fMRI-derived priors as constraints to demonstrate its ability to integrate multi-modality information.

### C. Benchmarks and performance metrics

The performance of Blk-Cham is compared to five benchmark ESI algorithms, including LORETA, EBI-Convex, ts-Cham, L21-Sissy and BESTIES:

- LORETA [6] is a widely-used linear method that applies the discrete Laplacian operator as constraints. The regularization parameter and can be estimated by maximizing the marginal likelihood P(*B*|*α*) [**?**].
- EBI-Convex [29] is a sparse Bayesian learning-based method, which utilizes the augmented lead-field for the joint estimation of both source activity and noise (refer to Eq. 1). Champagne algorithm is used to obtain the solution.
- ts-Cham [14] is a Bayesian framework that estimate source using two-stage champagne. In the first stage, original champagne is employed to find the potential locations of source. Spatial basis functions around the sources are subsequently constructed, and champagne is used again to estimate the final locations and extents of source based on the spatial basis functions.
- L21-Sissy [16] incorporates *L*_1_ constraints on both source and gradient domains to obtain sparsely structured source activities. Besides, *L*_21_ norm is imposed on temporal domain to ensure temporally continuous results.
- BESTIES [36] is a Bayesian approach that adopts firstorder Markov random field to model the spatial smoothness, and temporal basis functions (TBFs) to characterize temporal dynamics. The source is represented as a linear combination of TBFs and a temporal error term: *S* = *W* Φ + *E*.

To quantitatively assess the performance of the proposed methods, four validation metrics are employed: AUC (area under precision-recall curve [26]), that measures the sensitivity and specificity in reconstructing source locations and extends; DLE (distance of localization error), that estimates the localization error of reconstructed sources; MSE (relative mean square error), that measures the temporal error of estimated sources; and CORR (correlation), that measures the temporal accuracy of ESI results. The detailed computations of these metrics are provided in Appendices.

## IV. Results

We analyze the impact of SNR levels, source extents, and the number of sources, comparing Blk-Cham with five benchmark ESI algorithms. The effectiveness of different strategies for incorporating prior constraints is also assessed through simulations. Additionally, ESI results from three realistic datasets are presented to showcase the practical applicability of Blk-Cham. The validation results are detailed below.

### A. Results of simulation experiments

#### 1) Influence of SNR

Fig. 2 shows two representative examples of reconstructed sources obtained from different ESI methods under high (10 dB) and low (0 dB) SNR conditions. In terms of source extents, the LORETA and L21-Sissy yield spatially blurred results, while EBI-Convex and BESTIES produce overly focal sources. As for the reconstruction of time courses, we find that LORETA and L21-Sissy are sensitive to SNR, with reconstructed time courses significantly contaminated with noise at low SNR. In contrast, ts-Cham and Blk-Cham yield superior performance at both SNR levels. Moreover, the performance metrics across various SNR levels (i.e., −5, 0, 5, 10 dB) are presented in Fig. 3. As expected, all methods show improved performance in both spatial extent and temporal aspects with the increase of SNR. Moreover, the performance metrics demonstrate that Blk-Cham outperforms the benchmark methods in terms of robustness to noise, consistently providing accurate estimations of both spatial extents and temporal dynamics, even in challenging conditions

**Fig. 2.**
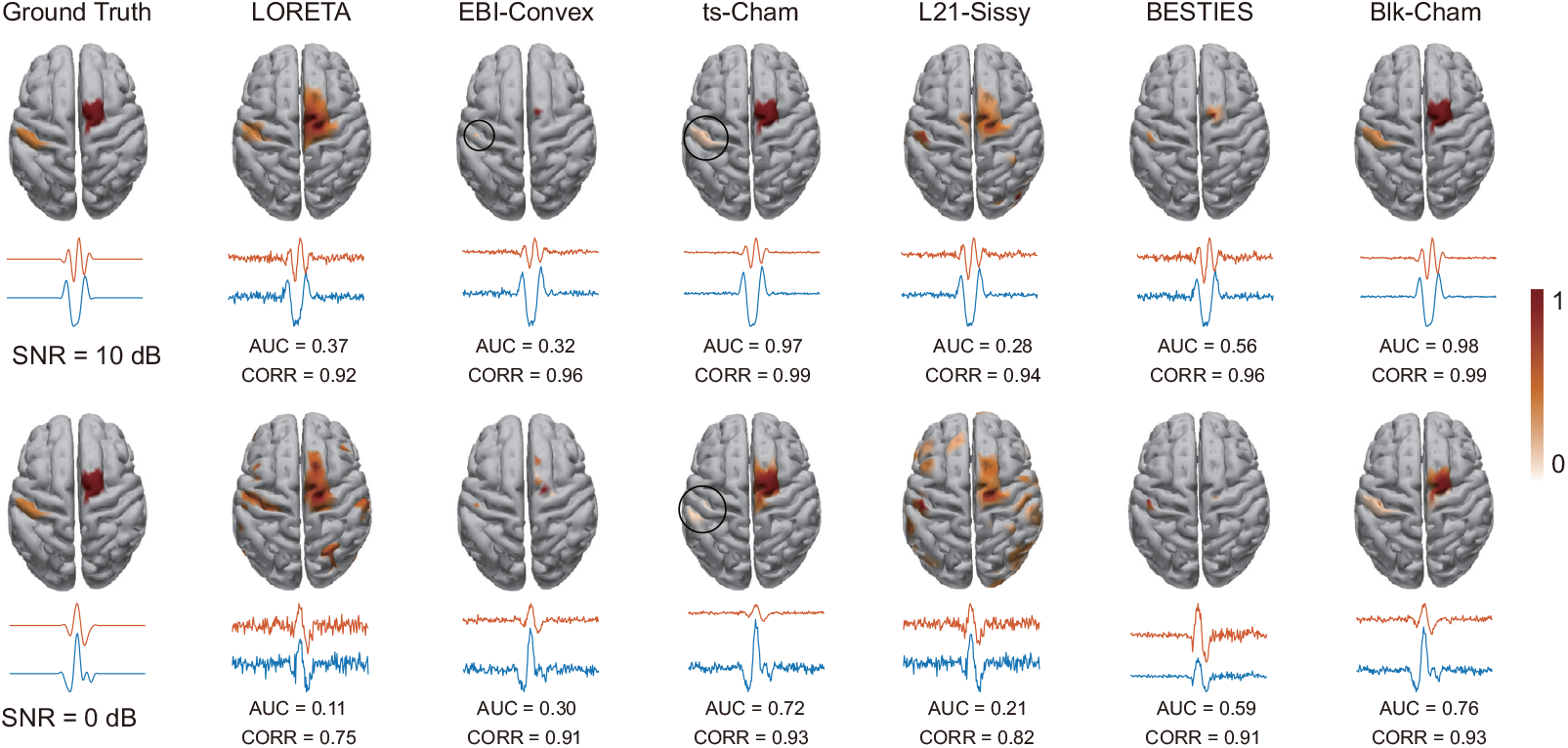
ESI results of different methods under different SNRs (i.e., 10 and 0 dB). Two sources are activated and the source extents are around 10 cm^***2***^. The source maps are presented as the power of estimated sources (normalized into 0 ***−*** 1). The reconstructed source time series are averaged within each source patch. The threshold is automatically determined by Otsu’ methods. Part of the reconstructed sources are encircled for better presentation purpose.

**Fig. 3.**
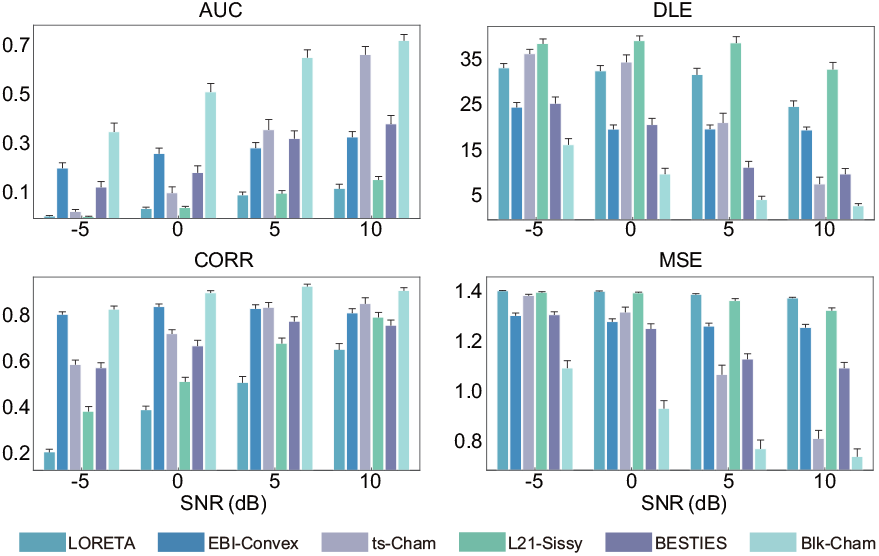
Performance metrics for ESI methods under various SNRs (i.e., −5, 0, 5, 10 dB). The source number is 2 and the extent of each source is around 5 cm^***2***^. The results are presented as mean ***±*** SEM (standard error of the mean).

#### 2) Influence of source extent

In Fig. 4, we show two representative examples of ESI results under small and relative large extents of a single patch of source. The LORETA produces over-smoothed source estimations, while EBI-Convex, BESTIES and L21-Sissy tend to yield focal results. EBI-Convex, which favors sparse solutions, performed best when the source extent was highly focal. In contrast, ts-Cham and Blk-Cham accurately reconstructed source extents in both cases. The performance metrics under various source extents are presented in Fig. 5. The proposed Blk-Cham outperforms the benchmark methods. It is noteworthy mentioning that when the source extent is small, i.e., 1 cm^2^, Blk-Cham slightly overestimates the extents due to its block-wise modeling of each dipole and its neighbors. In the simulation, each block contains about 7 dipoles (out of 4747 dipoles), while the focal source typically involves fewer than 3 dipoles. However, considering neurophysiological evidence that a minimal source extent is necessary for detectable E/MEG signals [4], [5], the block-wise approach of Blk-Cham is both reasonable and wellsuited for practical applications.

**Fig. 4.**
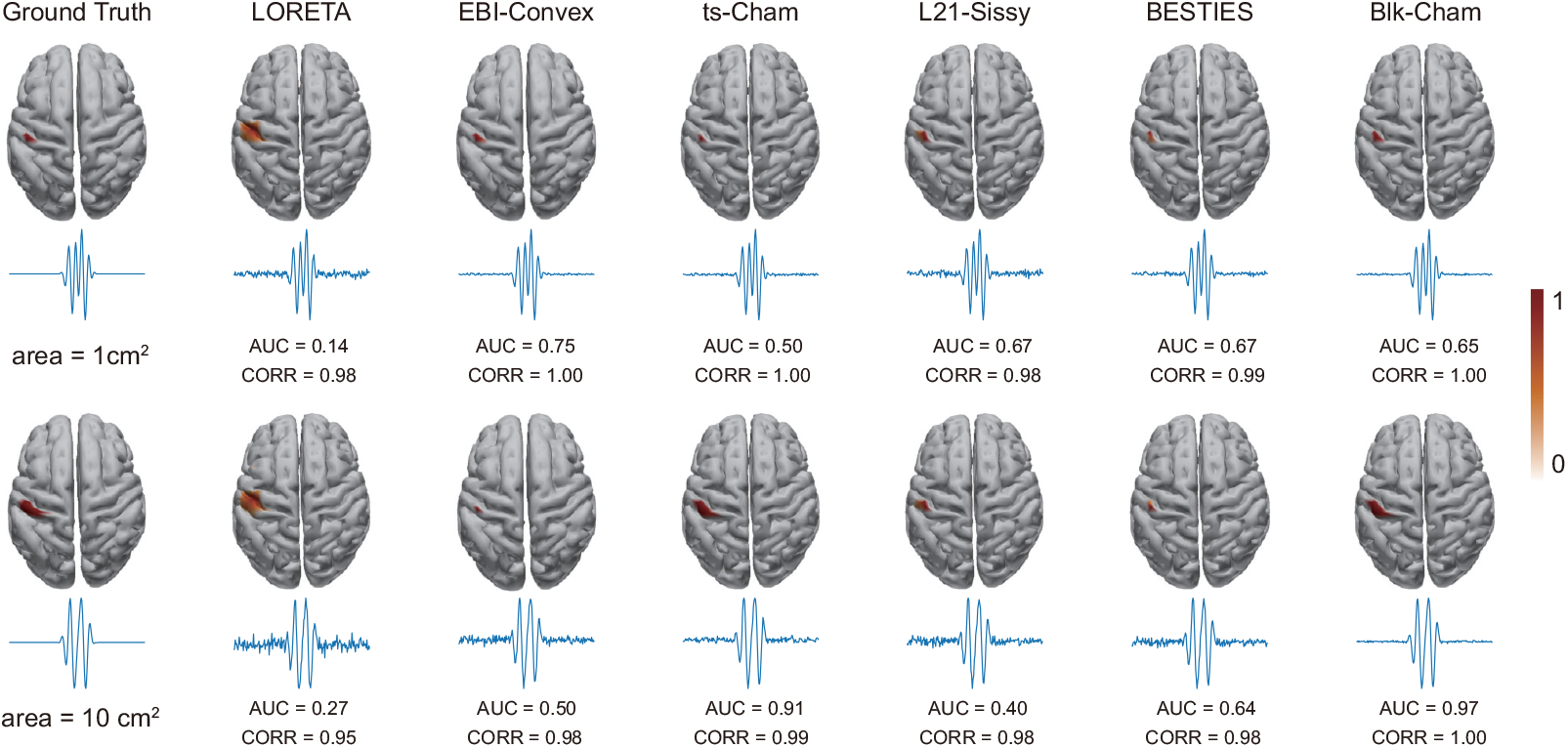
ESI results of different methods under different source extent (i.e., around 1 and 5 cm^***2***^). One sources are activated and the SNR is set to 10 dB. The source maps are presented as the power of estimated sources (normalized into 0− 1). The reconstructed source time series are averaged within each source patch. The threshold is automatically determined by Otsu’ methods.

**Fig. 5.**
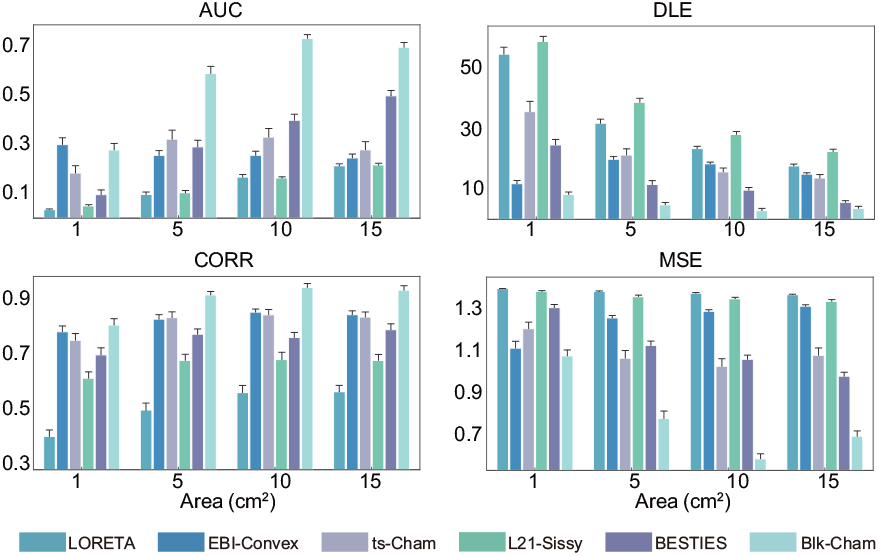
Performance metrics for ESI methods under various source extent (i.e., 1, 5, 10, 15 cm^***2***^). The source number is 2 and the SNR is 5 dB. The results are presented as mean ***±*** SEM (standard error of the mean).

#### 3) Influence of the number of source

Fig. 6 presents a representative example of source imaging results obtained from the 6 methods. LORETA and L21-Sissy fail to reconstruct source 3 while Blk-Cham accurately estimate the strength and extents of sources. The performance metrics under various numbers of source (i.e., 1, 2, 3, 4) are illustrated in Fig. 7. The Blk-Cham consistently outperforms the benchmark methods, which achieves superior performance in reconstructing both spatial and temporal aspects of source activities. Overall, the results highlight the robustness and superiority of the proposed Blk-Cham in accurately reconstructing source activities under varying source configurations.

**Fig. 6.**
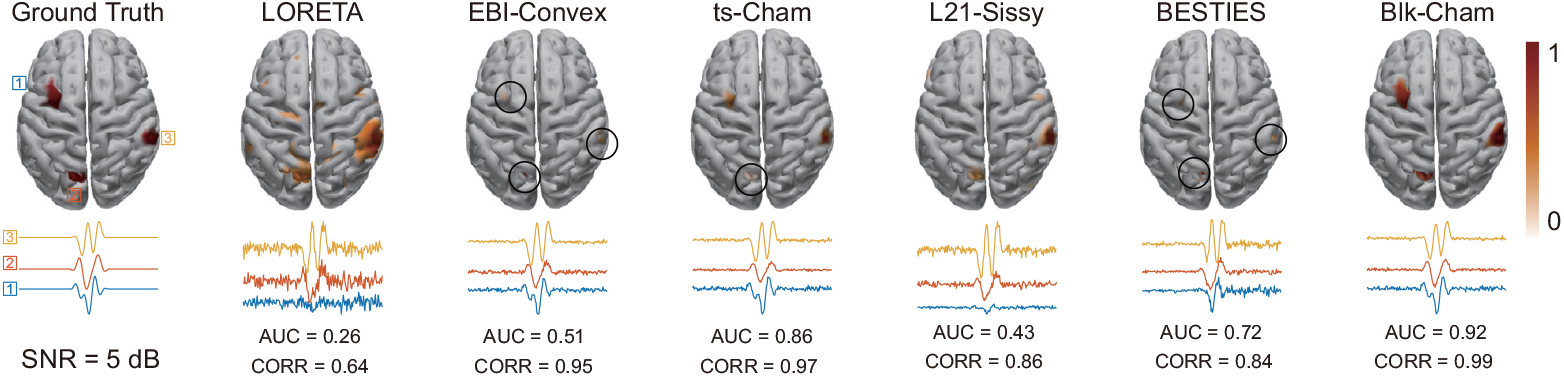
ESI results of different methods with three source patches. The source extents are around 5 cm^***2***^ and the SNR is set to 5 dB. The source maps are presented as the power of estimated sources (normalized into 0 − 1). The reconstructed source time series are averaged within each source patch. The threshold is automatically determined by Otsu’ methods. Part of the reconstructed sources are encircled for better presentation purpose.

**Fig. 7.**
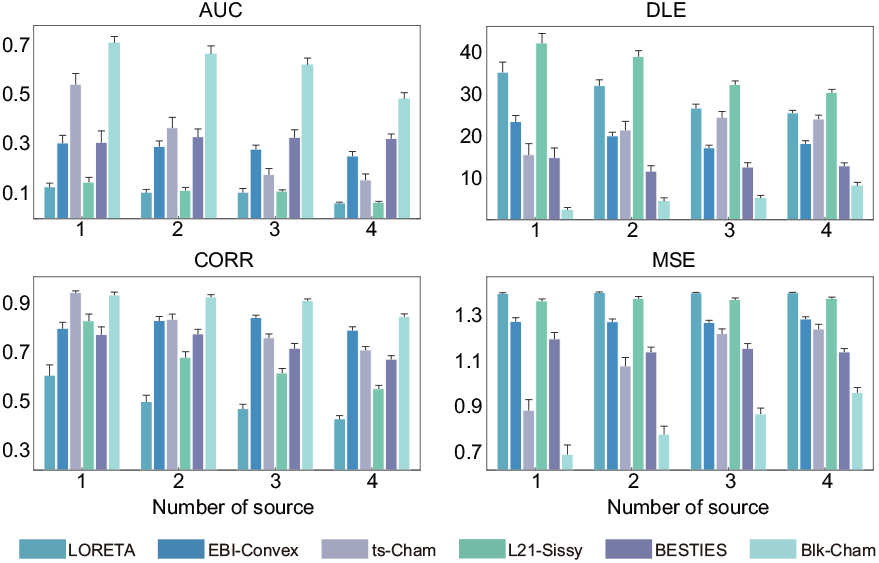
Performance metrics for ESI methods under different number of source (i.e., 1, 2, 3, and 4). The SNR is 5 dB and the extent of each soure patch is around 5 cm^***2***^. The source maps are presented as the power of estimated sources (normalized into 0***−***1). The results are presented as mean ***±*** SEM (standard error of the mean).

#### 4) Incorporation of prior knowledge

Fig. 8 presents an example of ESI results obtained from Blk-Cham incorporated with different prior constraints, as well as the performance metrics under different SNR levels: 1. exact prior, where the prior knowledge is exactly consistent with the ground truth; 2. scale prior, where the locations of prior knowledge aligns with the ground truth, while the extents are different, varying between 0 % to 50 % larger or smaller than the actual extent; 3. mixed prior, where the prior involves scale prior and 1 to 4 incorrect ones. we have scaled the values of DLE for illustration purpose: DLEn = 10*/*[− log(DLE) + 20]. The results demonstrate that both the exact and scale prior strategies significantly enhance the reconstruction of the spatial properties of the sources, particularly under low SNR conditions. As for the mixed prior strategy, we observed improved performance compared to the original method at low SNR cases. When the SNR is high, the original Blk-Cham can perform robustly without the need for additional priors.

**Fig. 8.**
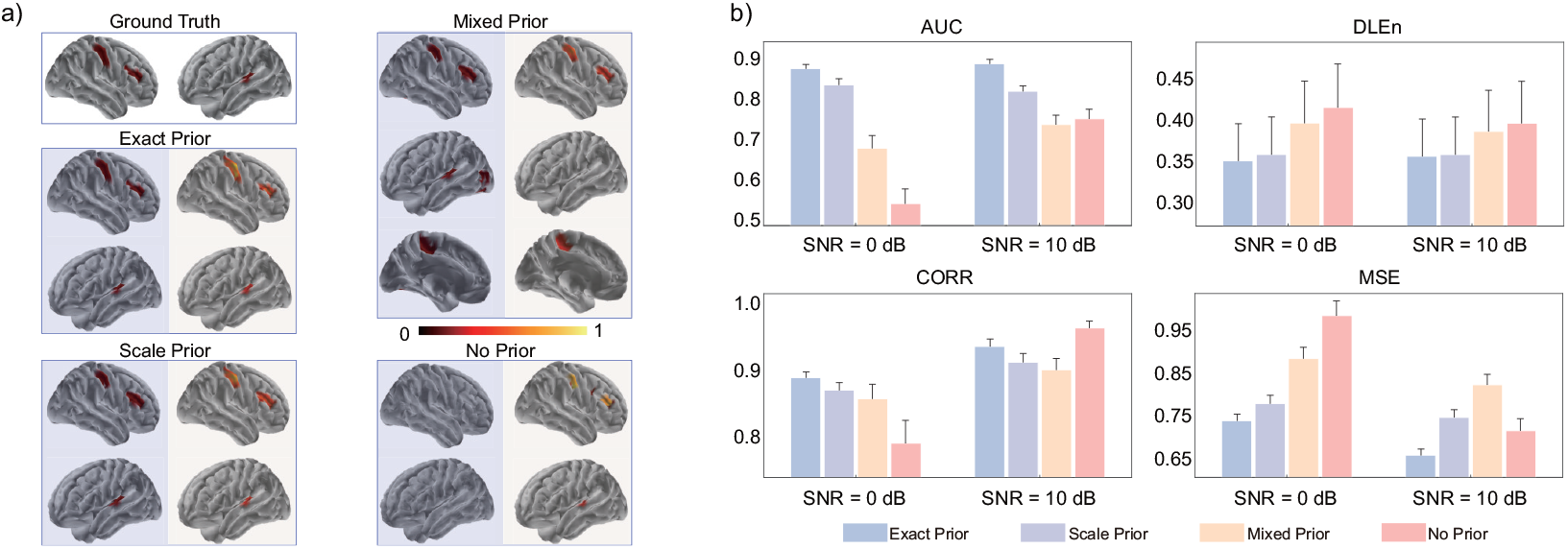
(a) ESI results of different strategies for incorporation priors. The source extents are around 10 cm^***2***^ and the SNR is set to 5 dB. The source maps are presented as the power of estimated sources (normalized into 0− 1). The threshold is automatically determined by Otsu’ methods. Exact prior: the prior knowledge is exactly consistent with the ground truth; Scale prior: the locations of prior knowledge are consistent with the ground truth, while the extents are different with the ground truth; Mixed prior: the prior involves these from scale prior and 1 to 4 incorrect ones. For each subgraph, the left panel shows the weights of prior constraints, and the right panel shows the ESI results. (b) Performance metrics under different SNR (i.e., 0 and 10 dB) with 2 source patches. The values of DLE are scaled for illustration purpose.

### B. Results of real Data

#### 1) localize-MI data

Fig. 9 presents the source imaging results of 2 subjects obtained from 5 ESI methods. For each subject,the ground truth of stimulation sites, i.e., the location of the pair of stimulating contacts (marked by red and yellow point), as well as their center (marked by blue point), is presented for intuitive assessment. We also highlight the location of ground truth with a green point in the ESI maps. For subject #1, LORETA, EBI-Convex and Blk-Cham successfully identify the stimulation sites in the right motor cortex. For subject #7, all the 5 methods reconstruct the generator sites in the right temporal lobe. In contrast, the proposed Blk-Cham not only successfully locates the stimulation sites for both subjects, but also yield structurally-sparse results that offer valuable insights into the source extents.

**Fig. 9.**
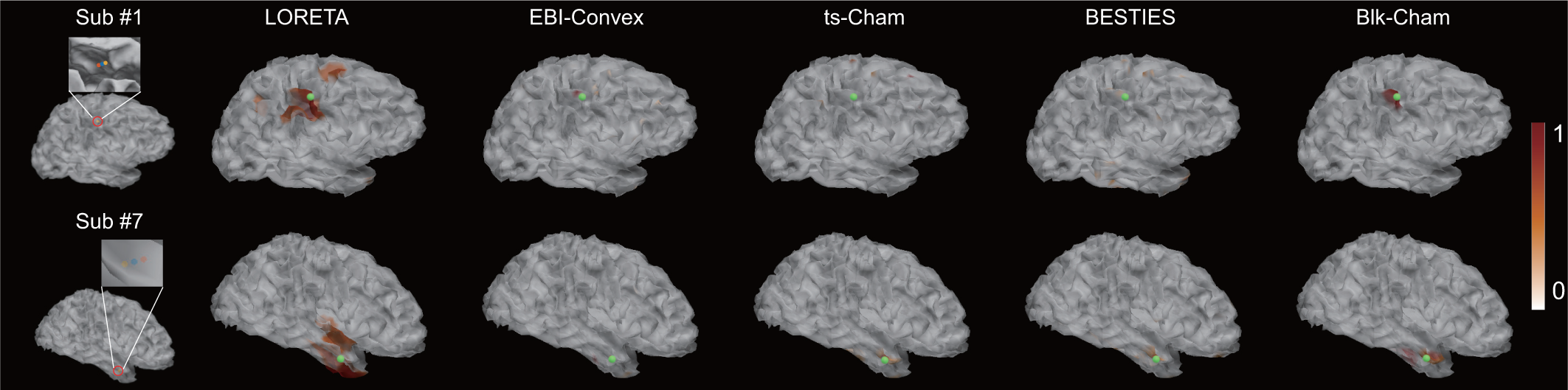
ESI results of deep brain stimulation obtained from LORETA, EBI-Convex, ts-Cham, BESTIES and Blk-Cham. The location of stimulation contacts (marked by red and yellow point) as well as the center of them (marked by blue point) are regarded as the ground truth of stimulation-induced activity. The location of ground truth is also highlighted in the ESI maps with green point.

#### 2) Epilepsy data

The source imaging results at the peak of averaged spike data are presented in Fig. 10. The results obtained from 4 methods are consistent with clinical findings that the seizure zone locates at the left frontal lobe [34]. The LORETA produces smooth sources, while the BESTIES and ts-Cham yield focal results. In contrast, Blk-Cham provides a clear boundary between activated and background sources, highlighting its ability to accurately estimate the extents of epilepsy-related brain regions. This demonstrates the efficacy of Blk-Cham in delineating source extents, which is crucial for identifying and characterizing the epileptic zone.

**Fig. 10.**
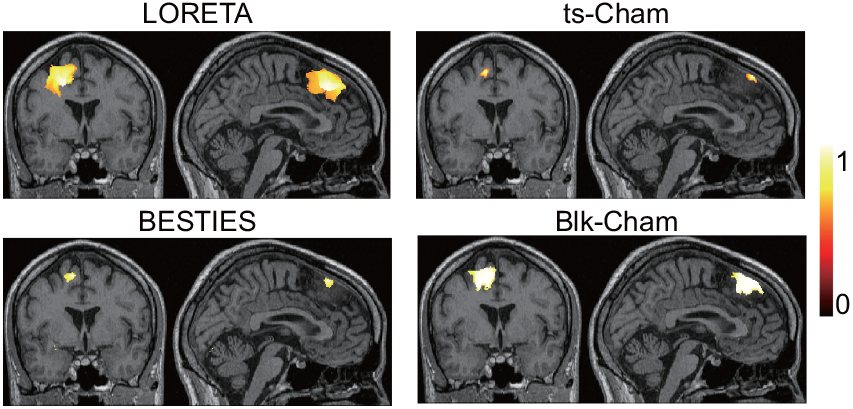
ESI results of epilepsy data obtained from LORETA, ts-Cham, BESTIES and Blk-Cham. The results are presented at the peak of spike.

#### 3) face-processing data

The ESI results of face-processing ERP are presented in (Fig. 11). According to the topographic analysis of ERP, fMRI responses, as well as our previous findings [37], the face-processing ERP data mainly involves two stages:1). 80 − 130 ms post-stimulus that locate at the bilateral visual cortex (corresponding to the P1 component); 2). 130 − 200 ms post-stimulus that locate at fusiform gyrus (corresponding to the N170 component). We apply LORETA, Blk-Cham and Blk-Cham with prior constraints derived from fMRI (Blk-Cham-prior) to reconstruct the underlying source activities. The results at T1 (100 ms post-stimulus) and T2 (170 ms post-stimulus) for each method are presented. The four approaches yield consistent source imaging results at both stages. Moreover, the proposed Blk-Cham with fMRI constraints could produce source estimations that align more closely with the fMRI response.

**Fig. 11.**
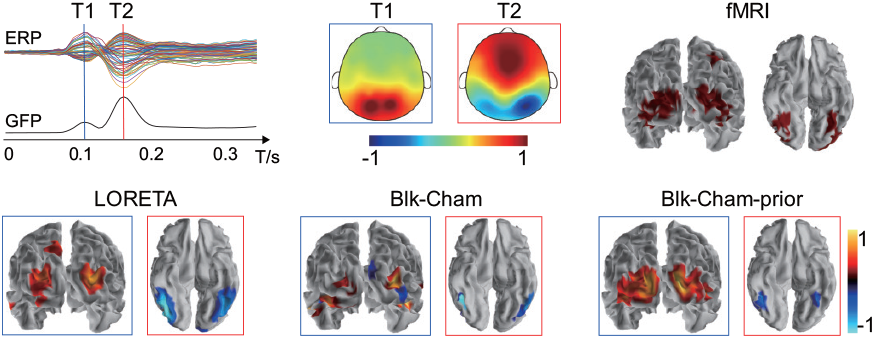
ESI results of face-processing ERP data obtained from LORETA, Blk-Cham and Blk-Cham-prior (i.e., Blk-Cham that incorporate prior constraints from fMRI). The topographies of ERP at T1 (100 ms post-stimulus) and T2 (170 ms post-stimulus), as well as task-related fMRI responses are presented. For each method, the source imaging results of ERP at T1 are highlighted with a blue box, and results at T2 are highlighted with a red box.

## V. Discussion

In this article, we propose a novel Bayesian framework, Blk-Cham, to accurately reconstruct the extended sources through integrating spatial information from local and remote scales. The Blk-Cham method leverages biophysical principles to model the local homogeneity of source activity, employing block-structured covariance that can be adaptively and automatically updated to improve the reconstruction of the source extents. Moreover, A key advantage of Blk-Cham is its ability to integrate multi-modality data, enabling the incorporation of remote spatial information as constraints to further enhance ESI performance. We conduct extensive validation through both simulated and realistic datasets to assess the feasibility and efficacy of Blk-Cham. These evaluations demonstrate that Blk-Cham outperforms benchmark methods in terms of accuracy and robustness, particularly in handling complex, extended sources in brain activity reconstruction.

Accurately estimating the extents of neural sources is of apparent importance in various ESI applications [10], i.e., identifying epileptic zones for planning subsequent interventions [17]. However, traditional methods based on *L*_2_-norm tend to overestimate the source extents, which requires a user-specific threshold to obtain meaningful results [3]. In contrast, the Bayesian learning framework provides an automated approach to identify activated sources, while it often produces sparse results scattered inside/around the ground truth with little information on the actual extents of sources [28]. To address this limitation, the Blk-Cham incorporates block-sparsity constraint under Bayesian framework, enabling automatic determination of relevant blocks to accurately reconstruct the source extents. Blk-Cham offers two key advantages over traditional SBF-based methods: 1). Blk-Cham models the block for each source and its neighbors, yielding overlapped blocks that can be flexibly combined to construct source with arbitrary extents. In contrast, traditional methods requires the construction of SBFs that match to the true areas of ground truth, which is often challenging. 2). While a fixed version of block yields satisfactory results, Blk-Cham also allows for data-driven update of blocks. The traditional SBF-based methods often rely on predetermined, non-updated blocks once they are constructed, which limits their adaptability. In contrast, the data-driven update fashion of Blk-Cham facilitate a more robust approach to source reconstruction. Additionally, another advanced strategy employs sparse constraint in the gradient domain, which promotes piecewise smooth solutions with sharp edges to reconstruct sources extents [17]. The Blk-Cham, on the other hand, operates in the original source domain, making it effective for integrating constraints from other modalities. Taken together, the proposed Blk-Cham provides a flexible and robust framework for accurately estimating source extents, addressing key challenges in traditional methods and advancing ESI applications.

Solving the ESI problem requires incorporating physiological knowledge to regularize the solution, but the properties of source activities remain incompletely understood. Thus, integrating prior knowledge from other modalities, particularly fMRI activation maps recorded concurrently with E/MEG, has been continuously explored to enhance ESI performance [20]. The proposed Blk-Cham offers an efficient strategy for incorporating additional constraints (e.g., derived from fMRI maps). Specifically, the Blk-Cham models source activities located within fMRI maps as a combination of local source activities and those from remote regions indicated by fMRI maps. Notably, Cai et al. [28] have adopted a similar strategy to reconstructing extended source under sparse Bayesian learning framework. However, their approach used the mean lead-field matrix of the corresponding remote regions to represent the remote activities, which may lead to a loss of spatial information. In contrast, Blk-Cham introduce a more flexible approach by modeling the fMRI-derived constraints as a symmetric covariance matrix between sources, which is subsequently updated in a data-driven manner. Given that the generation of E/MEG and fMRI signals are not identical [23], this strategy effectively balances prior knowledge with the evidence from data. Despite its robustness to some degree of incorrect prior knowledge, as demonstrated in our simulations, Blk-Cham’s performance can still be affected when the priors are invalid. Specifically, erroneous priors may degrade ESI performance both spatially—by introducing spurious sources—and temporally—by including noise activities from irrelevant sources. Therefore, we emphasize the importance of carefully constructing modality-derived constraints to ensure their validity and relevance for the ESI problem.

From a practical perspective, several factors must be considered for the successful implementation of Blk-Cham. Among these, the SNR plays a critical role in source estimation, as E/MEG signals typically exhibit low SNR. A common approach involves calculating noise covariance from baseline activities and whitening the signals before source imaging. However, obtaining reliable noise covariance is often challenging in real application scenarios. To address this limitation, the proposed Blk-Cham method incorporates an augmented leadfield strategy to model noise activity and learns the residual covariance directly from the data [38], [29]. This approach enhances Blk-Cham’s robustness to varying SNR levels, enabling effective source estimation even in the absence of accurate noise covariance information. In this study, we assume an independent prior distribution for noise activity. However, an improved strategy could be employed to learn the inherent structure of the noise covariance more effectively [39]. Besides, the selection of temporal periods of E/MEG signals may also affects the ESI results. While Blk-Cham is capable of handling signals of arbitrary length, including single snapshot cases, using an appropriate temporal window −such as one that captures the microstate dynamics of E/MEG [37] − can lead to optimal results in real-world applications. Finally, the blocks used in this model can be extended to include higherorder neighbors for each dipole. This extension may enhance the reconstruction of sources with larger extents since Blk-Cham inherently favors sparse solutions.

## VI. Appendix

### A. Calculation of performance metrics

Let *S* and *Ŝ* denote the simulated and estimated sources respectively, *q* = *diag*(*SS*^⊤^) and 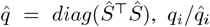 denote the energies of *i*^*th*^ element of simulated/estimated source.

#### 1) area under precision-recall (PR) curve (AUC)

the AUC is the area under PR curve, which is a plot of Recall over the Precision of the estimations as threshold *T*_*h*_ varies. For a specific *T*_*h*_, the dipole is assumed to be active if 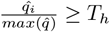.

#### 2) distance of localization error (DLE)

DLE is used to measure the localization error, which is defined as the average distance between the *k*^*th*^ dipole in the simulated map and the dipole with maximum energy among the 𝕀_*k*_:

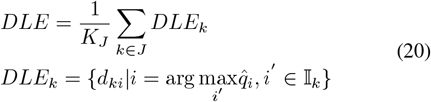

where *K*_*J*_ denotes the number of elements in J, which is the set containing the indices of the detected true sources. *DLE*_*k*_ measures the distance between the *k*^*th*^ true source and the estimated source with the maximum energy in 𝕀_*k*_, and *DLE* is the average of *DLE*_*k*_ over the true source indices *k*.

#### 3) relative mean square error (MSE)

MSE measures the reconstruction error between normalized time series of simulated and reconstructed time series: 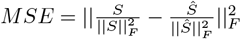

#### 4) correlation (Corr)

the Corr measures correlation (*corr*) of mean time courses in simulated source patches (*J*) of *S* and 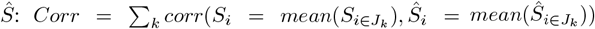 where *J*_*k*_ denotes the *k*^*th*^ simulated source patch.

## Footnotes

1 https://gin.g-node.org/ezemikulan/Localize-MI

2 https://neuroimage.usc.edu/brainstorm/Tutorials/Epilepsy

## Notes

This work was supported by the Zhejiang Provincial NSFC (LR23F010003) and by NSFC (82172056).

### Competing Interest Statement

The authors have declared no competing interest.

